# YAPP-CD: Yet another protein-peptide complex database

**DOI:** 10.1101/2021.06.16.448765

**Authors:** Joon-Sang Park

## Abstract

Protein-peptide interactions are of great interest to the research community not only because they serve as mediators in many protein-protein interactions but also because of the increasing demand for peptide-based pharmaceutical products. Protein-peptide docking is a major tool for studying protein-peptide interactions, and several docking methods are currently available. Among various protein-peptide docking algorithms, template-based approaches, which utilize known protein-peptide complexes or templates to predict a new one, have been shown to yield more reliable results than template-free methods in recent comparative research. To obtain reliable results with a template-based docking method, the template database must be comprehensive enough; that is, there must be similar templates of protein-peptide complexes to the protein and peptide being investigated. Thus, the template database must be updated to leverage recent advances in structural biology. However, the template database distributed with GalaxyPepDock, one of the most widely used peptide docking programs, is outdated, limiting the prediction quality of the method. Here, we present an up-to-date protein-peptide complex database called YAPP-CD, which can be directly plugged into the GalaxyPepDock binary package to improve GalaxyPepDock’s prediction quality by drawing on recent discoveries in structural biology. Experimental results show that YAPP-CD significantly improves GalaxyPepDock’s prediction quality, e.g., the average Ligand/Interface RMSD of a benchmark set is reduced from 7.60 Å/3.62 Å to 3.47 Å/1.71 Å.

## 1. Introduction

Recently, peptide docking has drawn more attention as protein-peptide interactions play key roles in many biological processes, and there is an increasing demand for peptide-based pharmaceutical products [1,2]. Several computational tools for peptide docking have become available, such as GalaxyPepDock [3], pepATTRACT [4], MDockPeP [5], CABS-dock [6] ClusPro [7], PIPER-FlexPepDock [8], HPEPDOCK [9], InterPep [10], GOLD [11] Surflex-Dock [12], DynaDock [13], HADDOCK [14], PEP-FOLD 3 [15], DINC [16], and AutoDock CrankPep [17]. They can be categorized into template-based and template-free docking methods. Template-based docking methods use known protein-peptide complexes as templates to generate predictions of unknown protein-peptide complexes, whereas template-free docking methods perform searches for good fits between the given proteins and peptides without using templates. As expected, the quality of the predictions made by template-based methods depends on the existence of complex structures similar to those of the protein and peptide being investigated; if the coverage of the template database is wide enough, template-based methods can yield reliable results [18]. For this type of methods, the template database must be updated regularly to leverage recent discoveries in structural biology.

GalaxyPepDock is one of the most widely used template-based docking tools. It is free, easy to use, and accessible via a dedicated web server or downloadable precompiled binaries from http://galaxy.seoklab.org/. Unfortunately, the template database included in the GalaxyPepDock binary package at present is based on an obsolete database called PepBind [19], which comprises structure coordinate files released before 2015. Approximately 10,000 new entries are appended to the Protein Data Bank (PDB) [20] annually, suggesting clear opportunities for the template database to expand significantly. By increasing the coverage of the template database, the quality of predictions made by GalaxyPepDock can be improved.

In this study, we present a protein-peptide complex database called Yet another protein-peptide complex database or YAPP-CD. It is made to be directly plugged into the GalaxyPepDock binary distribution and replace the native template database to improve GalaxyPepDock’s prediction quality. It consists of 19,533 protein-peptide complexes with standard residues collected from PDB. The experimental results show that using our new database significantly reduced the average Ligand/Interface root-mean-square deviation (RMSD) from 7.560 Å /3.620 Å to 3.469 Å /1.712 Å, respectively. The database is freely available to download at http://wnl.cs.hongik.ac.kr/yapp.

## 2. Materials and Methods

Several protein-peptide complex databases have been reported in the literature. LEADS-PEP [21], PepPro [22], and PeptiDB [23] are small sets (with 100 or fewer entries) mostly utilized in the benchmarking of docking tools. Larger databases with 1,300–3,200 complexes, such as PepX [24], PepBind [14], and PixelDB [25], have been proposed, but they are no longer maintained and are thus obsolete. The template database distributed with GalaxyPepDock’s binary package was based on PepBind. PepBDB [26] and Propedia [27] are recently released databases with over 13,000 complexes maintained to date. Some protein-peptide databases provide web interfaces for versatile functionalities, such as visualization and clustering, to help users understand protein-peptide interactions. In fact, protein-peptide complex datasets can also be derived from more comprehensive protein-ligand affinity databases such as BioLiP [28], because peptides can be considered a special type of ligand. BioLiP is comprised of 529,047 entries in total and 25,960 entries for peptide-type ligands (as of May 2021). Our proposed complex database YAPP-CD was built based on BioLiP. From the BioLiP annotation file, a list of complexes with peptide ligands was extracted, excluding peptides with non-standard residues, which resulted in 19,533 entries to be included in the database. Each complex entry consists of two coordinate files, one for the protein receptor and another for the peptide ligand, which are collected from BioLiP or PDB and manipulated to meet GalaxyPepDock’s requirements; that is, residues are renumbered such that the residue number always starts with one. Peptides with non-standard residues were excluded from our database because some of them caused GalaxyPepDock execution failures. We noticed that there were peptide ligands with non-standard residues included in GalalxyPepDock’s native template database; however, since GalaxyPepDock is distributed as a pre-compiled binary package, we were unable to debug and fix the problems with newly added non-standard residues. Therefore, we decided to exclude non-standard residue ligands from our database. In addition, it is unclear how non-standard residues contribute to the *interaction similarity* score-based template matching used in GalaxyPepDock. GalaxyPepDock generates complex structure predictions based on templates with the highest interaction similarity scores. Note that the *interaction similarity* score calculation makes use of the BLOSUM62 matrix, but it is not elucidated how residues that do not appear in the matrix**—**non-standard residues**—**are handled in the calculation [3].

GalaxyPepDock utilizes two kinds of molecular interactions found in each protein-peptide complex template to calculate the *interaction similarity* score: ionic and hydrophobic interactions. Information regarding the two types of molecular interactions for each protein-peptide complex is provided in our database. The interactions were identified using in-house Python scripts based on the following criteria: hydrophobic interactions are presumed if hydrophobic residues (namely, ALA, VAL, LEU, ILE, MET, PHE, TRP, PRO, and TYR) fall within the 5 Å range, and ionic residues, that is, ARG, LYS, HIS, ASP, and GLU, are considered to contribute to ionic interactions if they fall within the 6 Å range.

## 3. Results

In this section, we demonstrate the advantage of YAPP-CD by comparing the docking qualities of GalaxyPepDock with YAPP-CD and its native template database in terms of Ligand-RMSD, Interface-RMSD, and the fraction of native contacts (Fnat). LRMSD/IRMSD/Fnat calculation and prediction quality classification (e.g., *acceptable, incorret*) were performed based on the Critical Assessment of Predicted Interactions (CAPRI) criteria (http://www.ebi.ac.uk/msd-srv/capri/round28/round28.html).

As shown in Table 1, docking results for 56 PepPro [22] complexes demonstrated that YAPP-CD improved the prediction quality of GalaxyPepDock. By replacing GalaxyPepDock’s native template database with YAPP-CD, the average Ligand-RMSD/Interface-RMSD was reduced from 7.599 Å /3.618 Å to 3.469 Å /1.712 Å, respectively, and Fnat increased from 0.462 to 0.643. In addition, 47 (out of 56) complex structure predictions made by GalaxyPepDock with YAPP-CD could be classified as better than *acceptable* models according to the CAPRI classification, whereas only 30 predictions could be classified as better than *acceptable* models when docked with the native database. In other words, the number of successfully predicted complex structures using YAPP-CD was 17 more than the case without YAPP-CD. For instance, the best prediction made by GalaxyPepDock with YAPP-CD for the target PDB ID 5GTU showed an IRMSD of less than 2 Å and Fnat greater than 0.2, which is considered an *acceptable* model; however, the best predicted structure made without YAPP-CD exhibited a very high IRMSD of 18 Å and a very low Fnat of 0.04, which is considered an *incorrect* model. The other 16 targets successfully predicted only with YAPP-CD were 4YL6, 4QQI, 4M5S, 4K0U, 4J1V, 4HTP, 3WQ5, 3PLV, 3LU9, 3H8A, 2YBF, 2XS1, 2QOS, 2QN6, 2PEH, and 1WKW. In the docking experiments, 10 prediction models were generated by docking a specific peptide to the unbound structure and compared to the corresponding bound structure to calculate and find the best Ligand/Interface-RMSD for each target PDB ID listed in Table 1. (Details of peptide sequences are found in Appendix, Table A1.) In the PepPro benchmark database, 58 complexes were provided with their free form (or unbound) structures, but GalaxyPepDock failed to generate predictions with two targets, 5CQX and 3BEF, in our experiments. We used 2X0C instead of 5IC0 as the unbound structure for 5FZT because 2X0C is referred to as the free form of 5FZT in [29]. Note that 5IC0 is a triple-domain talin, whereas 5FZT and 2X0C are two-domain talin structures.

**Table 1.**
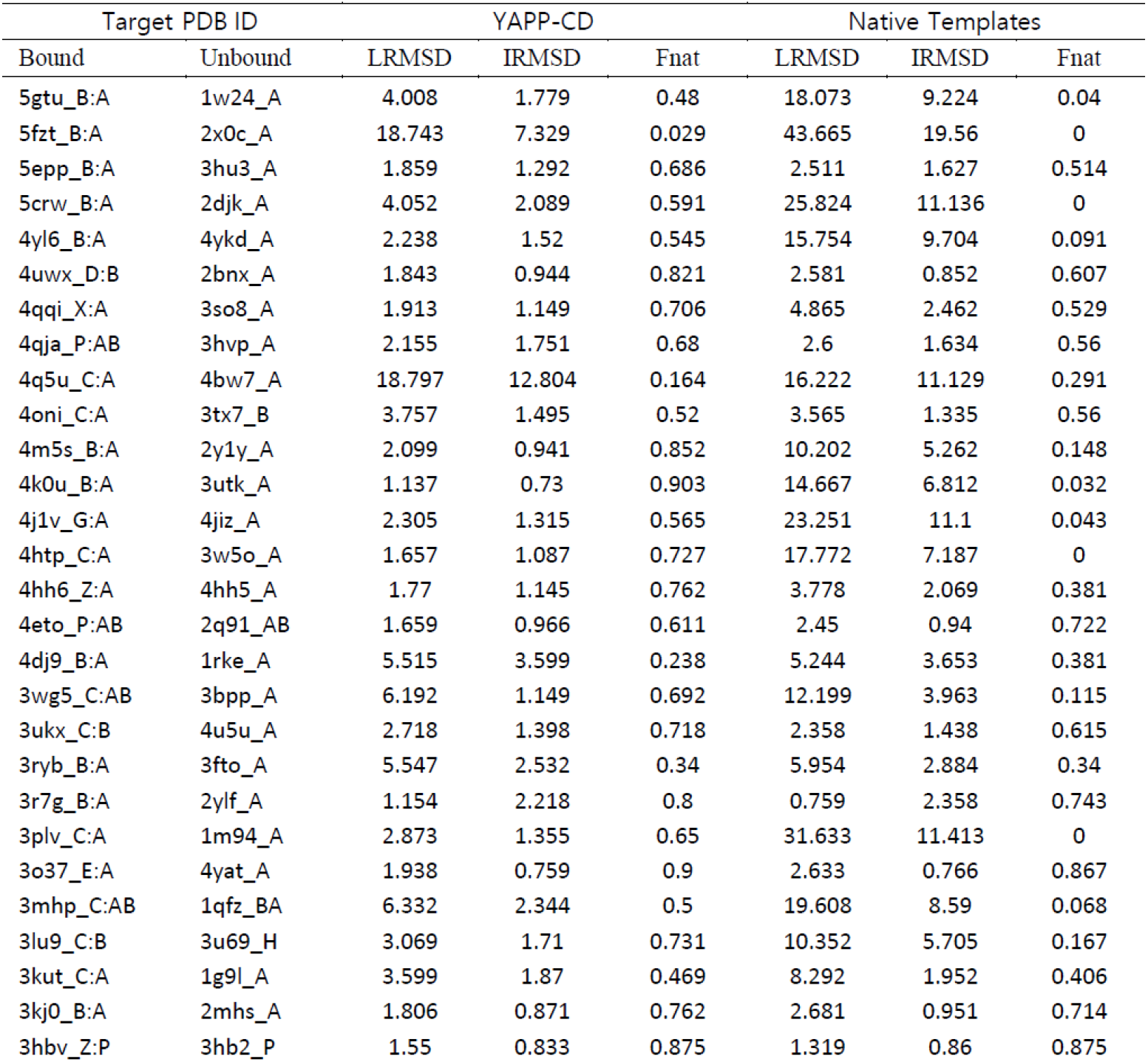
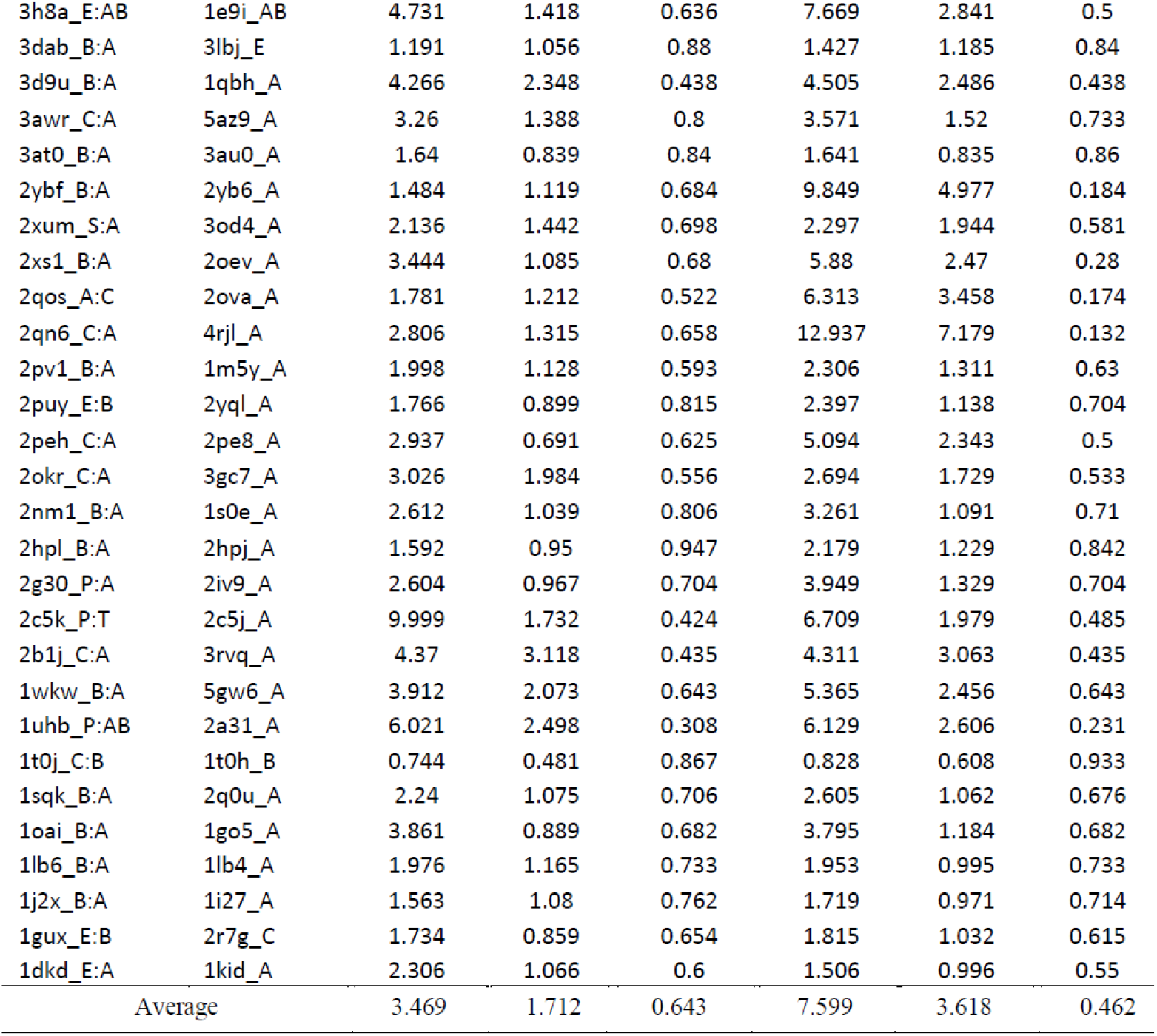
Docking results of 56 PepPro complexes. (For Bound structures, each PDB target is indicated as PDB ID followed by the peptide chain ID and its receptor chain ID. For Unbound structures, each PDB ID is followed by the receptor chain ID only.)

## 4. Discussion

Protein-peptide docking is a major tool for studying protein-peptide interactions, and several docking methods are currently available. Among the protein-peptide docking methods, template-based approaches such as GalaxyPepDock have been shown to yield more reliable results than template-free methods in recent studies. To obtain reliable results with a template-based docking method, the template database must be comprehensive enough and up-to-date to leverage recent advances in structural biology. In this paper, we present a protein-peptide complex database YAPP-CD, which is freely available at http://wnl.cs.hongik.ac.kr/yapp. YAPP-CD can be directly plugged into GalaxyPepDock to improve its prediction quality. Experimental results on a benchmark set of 56 targets showed that YAPP-CD improved GalaxyPepDock’s prediction quality, e.g., average Ligand/Interface-RMSD was reduced from 7.60 Å /3.62 Å to 3.47 Å /1.71 Å. At present, complexes with non-standard residue-based ligand peptides are not included in YAPP-CD; however, there are numerous complexes with non-standard residues in PDB, and some of them are believed to serve as meaningful templates for predictions. Our immediate next step toward expanding YAPP-CD is to develop strategies for including peptides with non-standard residues in the database.

## Appendix A

**Table A1.**
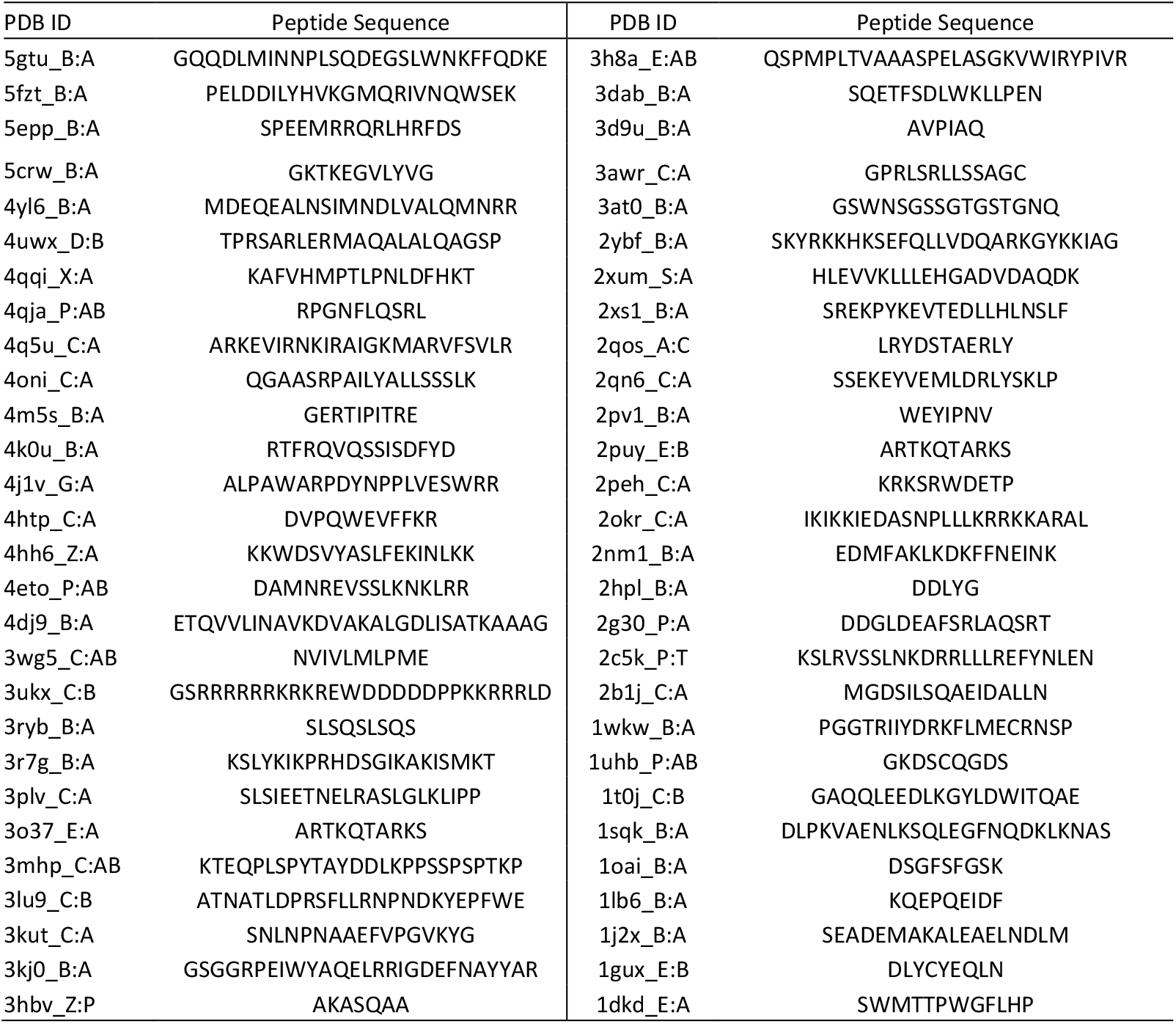
Peptide sequences of 56 PepPro benchmark structures used in docking experiments.

## Notes

### Competing Interest Statement

The authors have declared no competing interest.

